# A trait-based understanding of the vulnerability of a paleotropical moth community to predation by a sympatric bat with flexible foraging strategies

**DOI:** 10.1101/2023.08.10.552891

**Authors:** Pritha Dey, Rohini Balakrishnan

## Abstract

1. Prey profitability for a predator hunting diverse prey varies with species and seasons. Whereas prey selection by aerial-hawking or gleaning bats is well established, this is challenging to establish in species that adopt both these strategies flexibly. Measurable prey traits coupled with availability in the foraging grounds help characterize the vulnerability of prey species to predation.
2. In the Western Ghats of India, a global biodiversity hotspot, we studied an anthropogenic landscape, where insectivorous bats are abundant and diverse, but their impact on moth communities is little understood. We investigated the morphological traits of a sympatric moth community that make them more vulnerable to predation by *Megaderma spasma*, a bat with flexible foraging strategies. We also established the seasonal composition of moth prey in the diet of the bat.
3. We analyzed the discarded prey remains from several roosts, collected over three years, for seasonal patterns in the diet and selective hunting. Through light-trapping, we collected moth specimens in different seasons to assess the morphological traits of the moth community available in the foraging area of the bat.
4. The traits likely to affect the profitability of prey moths were measured: forewing length, hindwing length, wingspan, and body length (a proxy for body size); forewing area, hindwing area, maneuverability, and wing loading (as a proxy for evasive flight capability), and forewing aspect ratio (as a proxy for wing shape).
5. Our results showed that consumed moth prey diversity varies seasonally, with moths belonging to the Hepialidae family being preferred in the wet season. Moths belonging to the Sphingidae family were the most abundant in the diet, followed by Erebidae and Hepialidae. Sphingid moths have the lowest maneuverability, and highest forewing aspect ratio; the Hepialidae moths have the maximum body size followed by Sphingids thus confirming our hypothesis that larger moths with low evasive capabilities are more vulnerable to predation.
6. Assessing vulnerability at the prey community level, we establish a framework for future research on moth-bat interactions from the diverse and less-explored paleotropical communities. Additionally, the study reiterates the usefulness of trait-based approaches to understanding prey-predator dynamics.

## Introduction

Predator–prey relationships are central to community dynamics. The relationship involves strategies that predators adopt to become efficient hunters and those that prey adopt to avoid getting eaten (Schmitz, 2017). Recent studies have investigated these interactions as an evolutionary battle through functional traits which are measurable morphological, behavioural, physiological, or life history traits of organisms associated with their ecological functions (Ferry-Graham, 2002; Violle et al., 2007). Functional prey-predator relationships are based on the presumption that prey characteristics rather than prey taxonomy should drive prey selection by predators (Spitz et al., 2014).

Morphological traits related to flight efficiency and maneuverability help characterise the prey’s profitability to a predator based on how difficult it is to catch them or how much effort goes into searching for them. For example, hunting prey that are too large is energetically costly to predators and too small a prey is not worth the chase (Schmitz, 2017). Selecting a prey based on size is quite common (Brose, 2010; Nakazawa, 2017; Ortiz & Arim, 2016), but in addition, prey evasive tactics determine predator preference and its impact on the prey assemblage (Klecka & Boukal, 2013). Therefore, this trait-based approach is an important connection between evolutionary and community ecology that allows ecologists to study predator-prey dynamics at the scale of a community rather than a simple paired prey-predator model (Moore & Biewener, 2015; Schmitz et al., 2015; Schmitz & Trussell, 2016).

For 65 million years, moths and bats have been engaged in aerial warfare replete with stealth and deception (reviewed in ter Hofstede & Ratcliffe, 2016). As highly skilled and acrobatic hunters, bats choose prey that are energetically profitable and suited to their hunting skills (Arrizabalaga-Escudero et al., 2018; Mata et al., 2016; Vesterinen et al., 2016). Successful foraging by bats depends on the age and sex of the individual (Arizzabalaga et al., 2019; Mata et al., 2016), prey abundance (Wray et al., 2021), and prey accessibility due to habitat structure (Almenar et al., 2013). Selection of larger prey has been observed in insectivorous bats (reviews in Almenar et al., 2008; Burles et al., 2008; Fenton et al., 1990; Jones, 1990; Siemers & Schnitzler, 2000; Vesterinen et al., 2016) and attributed to prey profitability (energy gained/prey handling time) (Catania & Remple, 2005). Arrizabalaga-Escudero et al. (2019) provided a better understanding of prey resources using trait-based analyses, establishing that in a temperate ecosystem, a combination of functional traits of moths (wing loading and body mass) and life-history traits of bats (age and sex) governs predation risk from the insectivorous horseshoe bat *Rhinolophus euryale*. Fundamental to prey-predator interactions, these hypotheses are untested from the diverse tropical systems and from bats that follow different hunting strategies, such as megadermatid gleaning bats. Tropical arthropod communities are extremely high in species richness (Basset et al., 2012) which gives predators much to choose from and also allows them to specialize on profitable moths of certain species or traits.

Prey species have to be faster and have high maneuverability to escape predators successfully (Howland, 1974) and evasive flight is a common anti-predator tactic known in moths (Roeder, 1964, 1967). They “turn away” (directional flight) and/or “dive” (erratic flight) toward the ground in response to a distant bat or close bat respectively (Roeder, 1962). Depending on their auditory capabilities (Minnaar et al., 2015; Schoeman & Jacobs, 2011) and size (Surlykke et al., 1999), moths show species-specific strategies that are a functional consequence of the morphological differences that contribute to maneuverability (Hügel & Goerlitz, 2019). It is known that the hindwings in the flapping flight of moths and butterflies are more important for take-off, acceleration, and maneuverability than for sustained flight (Jantzen & Eisner, 2008; Stylman et al., 2020). However, it is yet to be investigated how different morphological traits make moths of different sizes vulnerable to predation, especially by bats with flexible foraging strategies.

Our focal predator is the lesser false vampire bat, *Megaderma spasma*, found mostly across south-east Asia, NE India, and in the Western Ghats. The insect order Lepidoptera (mostly moths) forms about 10-12% of the diet of *M. spasma* (Davison & Zubaid, 1992; Raghuram et al., 2015) and does not vary across seasons (Prakash, 2020). Usually, moth species with audible wing beats and nocturnal flight (e.g Sphingidae, Saturniidae) are abundant in the diet (Balete, 2010). *Megaderma spasma* hunt within a kilometer of their daytime roost (Prakash et al., 2021) and return to the roost with prey, thereby providing culled remains through which the prey species can be identified (Brosset, 1962). Their large ears make them capable of detecting low-frequency, directional sounds due to prey movement (Wang & Müller, 2009). However, *M. spasma* most likely uses multiple foraging tactics as their short, broadband, multi-harmonic, frequency-modulated calls are more suited to catching prey by perch hunting (‘flycatching’), in high clutter habitats rather than surface gleaning (Tyrell, 1990).

The predator is well-studied in our study area and its roost locations have been mapped, from which we reliably know that it feeds mostly on orthopteran prey (Prakash et al., 2021; Raghuram et al., 2015). Given its foraging tactics and size, it is capable of handling larger prey. Lepidoptera (especially moths) is the most diverse insect group that it feeds on. It remains to be known what kind of moths are chosen, in which season, and what traits drive this decision.

In this paper, our goal was to investigate, in a paleotropical ecosystem, the following questions: 1) Is there a seasonal difference in the diversity of moth species hunted by *M. spasma ?* and 2) Which morphological traits of a sympatric moth community make them more vulnerable to predation? We predicted that the consumed moth prey diversity would change with seasons and that larger moths with low evasive capabilities would be preferred by *M. spasma*. Overall, our study aims to bridge the gap in prey-predator interaction research from the tropics with a community-level trait-based analysis to understand the functional demands on the moth assemblages imposed by a sympatric bat species. This study establishes a model prey-predator system for further exploring the bat-moth arms race from this part of the globe.

## Materials and Methods

### Study Area

The study was conducted in the human-modified landscape just outside Kudremukh National Park, in and around Kadari Village, Udupi district, Karnataka, India (13°21′N–75°08′E) in the Western Ghats, a global biodiversity hotspot (Fig. 1). The annual rainfall ranges from 600-800 cm, with the maximum rain from June to September. The maximum temperature varies from 21-34°C (April to July) and the minimum temperature from 12-18 °C (January to May) (Nagaraja et al., 2005). The Western Ghats, a 1600 km long chain of mountains along the west coast of India, are unique because of their geographical location, stable geological history, heavy rainfall, and good soil conditions, crucial to maintaining a variety of tropical forest ecosystems (Pascal et al., 2004). The natural vegetation (below 300 m), i.e our core study area, is semi-evergreen and human-influenced (Subramanian et al., 2005). The Areca nut (*Areca catechu*), Rubber (*Hevea brasiliensis*) plantations, and agricultural lands have replaced the natural forests. *Megaderma spasma* roosts predominantly in human habitation in this landscape and the roosts are regularly monitored.

**Figure 1.**
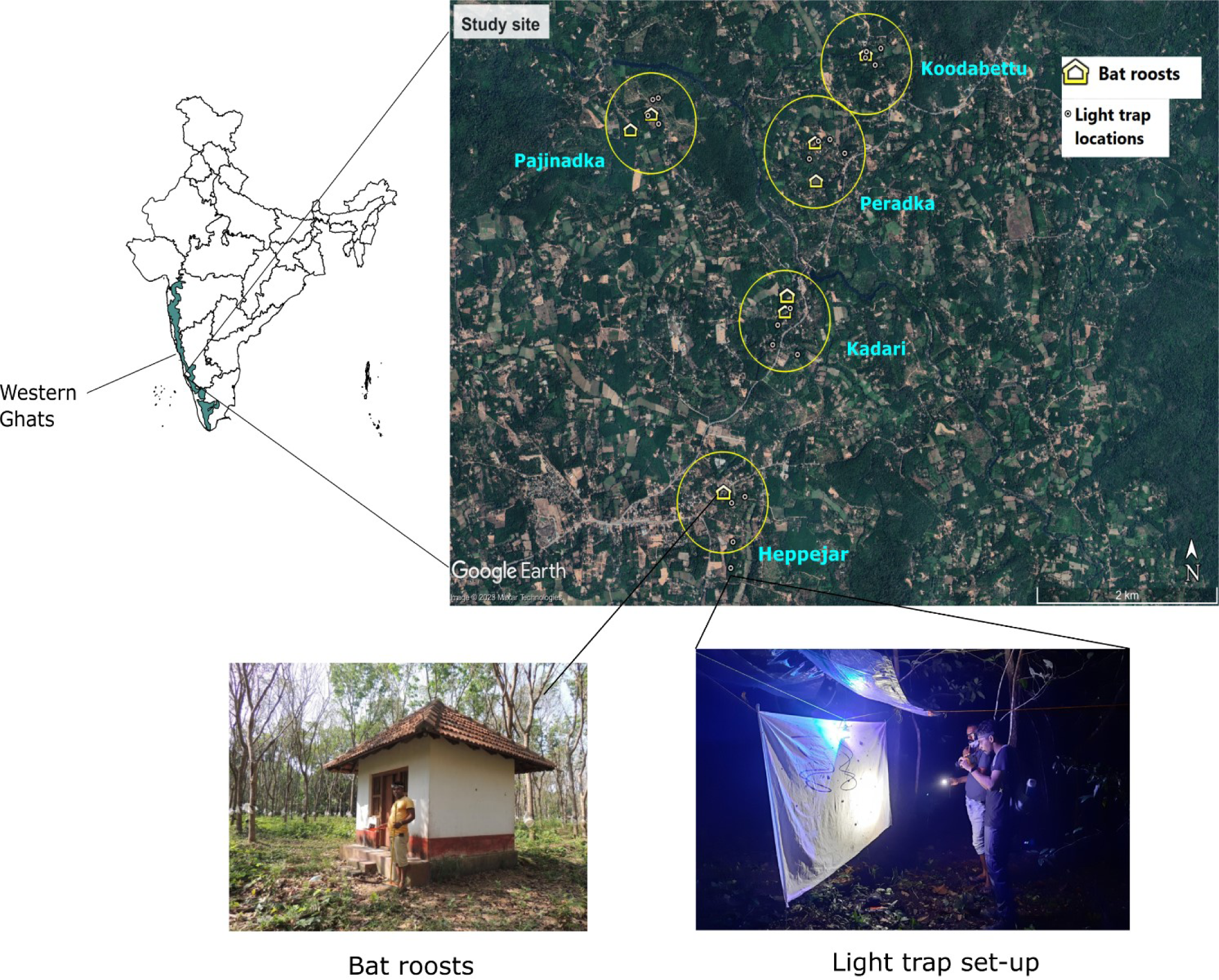
The map shows the location of the study area with details of the light traps around the bat roosts

### Culled remains collection

Prey remains discarded by bats on the floor of the roosts (referred to as ‘culled remains’) were collected weekly (2018-20, 2021) or every two weeks (2016-17). The collection was made from 21, 24, and 21 roosts in 2016-17, 2018-20, and 2021 respectively (details in Table 1 and Fig 1 in Supporting information). Some of the roosts become unavailable for collection for some parts of the year as they are private properties or they get damaged due to heavy rainfall. The culled remains collection was restricted to 8 selected roosts during the month of May-July 2021, due to a nationwide lockdown amidst the global COVID-19 pandemic, so the comparisons are done only among those 8 roosts for the wet season in the third year.

Most insect parts were classified to the order level, but the moth wing remains were identified to species level, whenever possible. We also identified each forewing as a left or right wing. By matching the left and right wings, we arrived at the minimum number of individuals for all insect orders. A database was created with segregated insect orders and the identified moth species over the years. About 40% of the culled remains could not be identified, as they were torn, or damaged due to rain or the wing patterns were not visible due to loss of scales.

### Light trapping

We used light traps to study moths at random points around 8 bat roosts at five sites (Field Station, Peradka, Pajinadka, Koodabettu, and Heppejar) (Fig. 1) for documenting the available moth diversity. The light trapping was done by the vertical sheet method (a white sheet measuring ∼90 x 190 cm), illuminated by LepiLED (Brehm, 2017) (Fig.1). LepiLED consists of power LEDs with peaks at 368 nm (ultraviolet), 450 nm (blue), 530 nm (green), and 550 nm (cool white), corresponding to the peak sensitivity of most Lepidoptera eye receptors (Brehm, 2017).

Light-trapping was done for 10 nights/month (each site was sampled twice a month) for 7 months from the fading moon to the new moon cycle (Jan-April; Aug-Oct) in 2021. We sampled moths with two traps running for different periods of the night because the storage life of portable batteries was a maximum of only 5 hours (details in Table 2 and Fig. 2 in the Supporting Information). The traps were placed within a radius of 1 km from a bat roost. The effective attraction distance of moths to light traps used in our study is about 50 m; thus, we spaced our traps by 100 m to minimize bias and maximize efficiency within a night at a sampling site.

**Figure 2.**
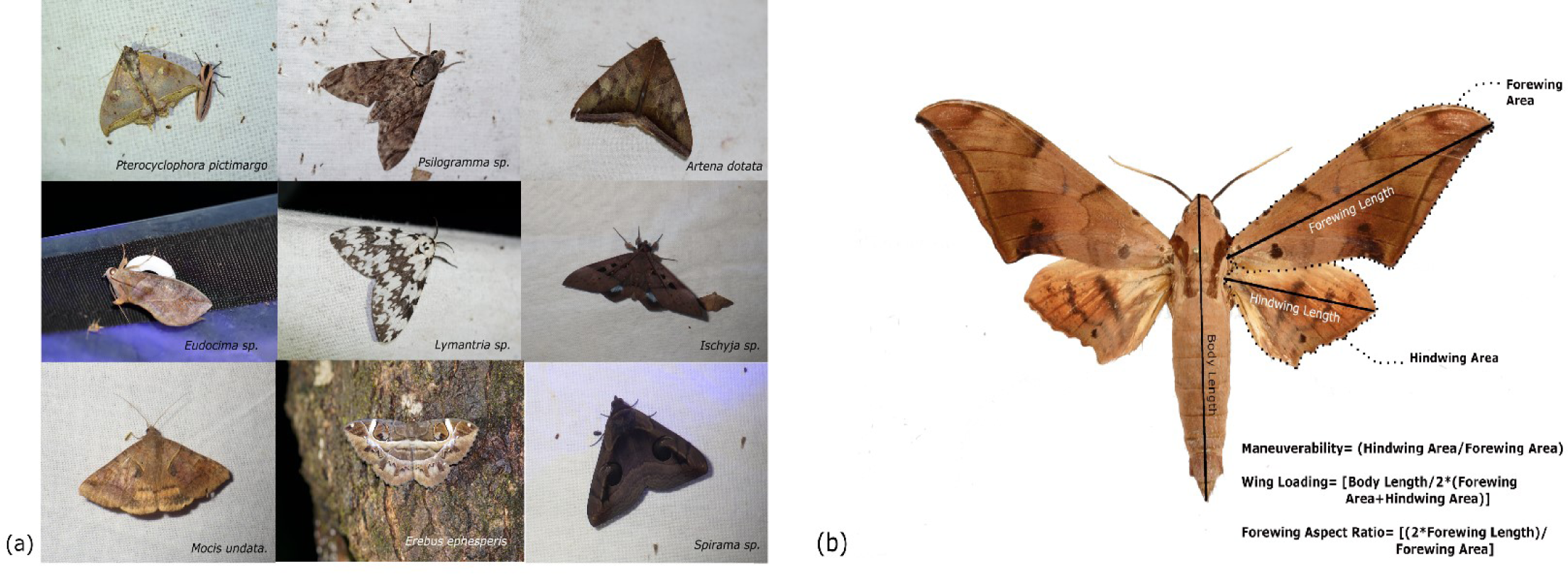
a) Representative moth species that were collected from the light traps and b) the morphometric traits measured from voucher specimens that were collected from the light traps and stored in the NCBS research collection

The number of individuals of moth species having >40 mm wingspan were counted every hour, as the smallest moth species identified from the culled remains collection of previous years had around 40 mm wingspan.

### Measuring morphological traits

The specimens collected from the light traps were deposited as voucher specimens at The Research and Collections Facility of the National Centre for Biological Sciences, Bangalore, India. Fig. 2a shows a representative moth species diversity of the light traps. We photographed the voucher specimens with their wings outstretched and with a reference scale. From these photographs, we measured the following traits using the software ImageJ (Schneider et al., 2012): Forewing length (FWL); Hindwing length (HWL); Body length (BL) (a proxy for body size); Wingspan; Forewing area (FWA); Hindwing area (HWA); Maneuverability=HWA/FWA (as a proxy for evasive flight capability, higher value equals higher capability) (Arrizabalaga-Escudero et al., 2019; Jantzen & Eisner, 2008); Wing loading =BL/2(FWA+HWA) and Forewing aspect ratio = (2*FWL)/FWA (as a proxy for wing shape) (Shi et al., 2015) (Fig. 2b). These traits were chosen as they are likely to affect the profitability of prey moths for *M. spasma*.

Only specimens having wingspan >40 mm were considered for the analyses. Each trait was measured from 2 to 5 specimens per species to obtain an average value. We had only one individual from the Family Crambidae, which was not included in the analyses. A total of 418 specimens of 158 morphospecies belonging to 14 families were measured.

### Replication Statement

| Section | Scale of inference | Scale at which the factor of interest is applied | Number of replicates at the appropriate scale |
| --- | --- | --- | --- |
| Culled remains collection | Roost | Year | 21, 24, and 21 roosts in 2016-17, 2018-20, and 2021 respectively |
| Light trapping | Location | Location | 4, 4, 4, 4, 4 (for each of the 7 months) |
| Morphometry | Specimen | Family | 418 specimens of 158 morphospecies in 14 families |

### Statistical Analyses

We used the light-trapping data as a proxy of the moth species pool present in the study area, and compared species found in the culled remains with those not found in the culled remains for further analyses. To understand sample completeness, we computed rarefied and extrapolated species accumulation curves for all the five sites sampled, using the iNEXT package (Hsieh et al., 2016) in R.

Culled remains: The diversity of moth species (Shannon Diversity Index) and the proportions of moth individuals belonging to different families found in the culled remains were calculated across seasons and for three years. As the proportion of individuals belonging to the three families Sphingidae, Erebidae (in all seasons), and Hepialidae (in the wet season) were higher, we compared the proportions of individuals in these families across seasons using a two-proportions z-test.

Morphological traits: All 9 morphological traits were compared among the species present and absent in the diet using a Wilcoxon Rank Sum Test. To compare the morphological trait space of the three most abundant families present or absent in the diet (Erebidae, Sphingidae, Hepialidae), we carried out a Principal Coordinates Analysis (PCoA) followed by PERMANOVA (Permutational multivariate analysis of variance) on a Euclidean distance matrix. The proportion of species present in the diet from the other families was low and thus was not considered for comparing the trait space. We also performed a multivariate beta diversity analysis to test for the homogeneity of dispersion within the trait space of each of the families. Both analyses were done with the *betadisper* and *adonis* functions in the Vegan package (Oksanen et al., 2009) in R.

To understand the effect of different morphological traits on the selection of prey by *M. spasma*, we performed logistic linear regression (binomial function) with the presence or absence in the diet as the response variable and the Body length, Maneuverability, Forewing Aspect Ratio, and Wing Loading as the predictors. Predictor variables that were correlated above 0.8 were not considered, to avoid multicollinearity in the models (Fig. 3 in Supplementary material). All statistical analyses and visualizations were performed using R (“R Development Core Team: R: A language and environment for statistical computing,” 2008).

## Results

### Species pool at the foraging grounds

We carried out light-trapping for 7 months in 2021, to establish the prey moth species pool available for *M. spasma* around their roosts. Our sampling was adequate at the five sampling sites based on the abundance-based sample completeness curve of species sampled at each site (Fig. 3).

**Figure 3:**
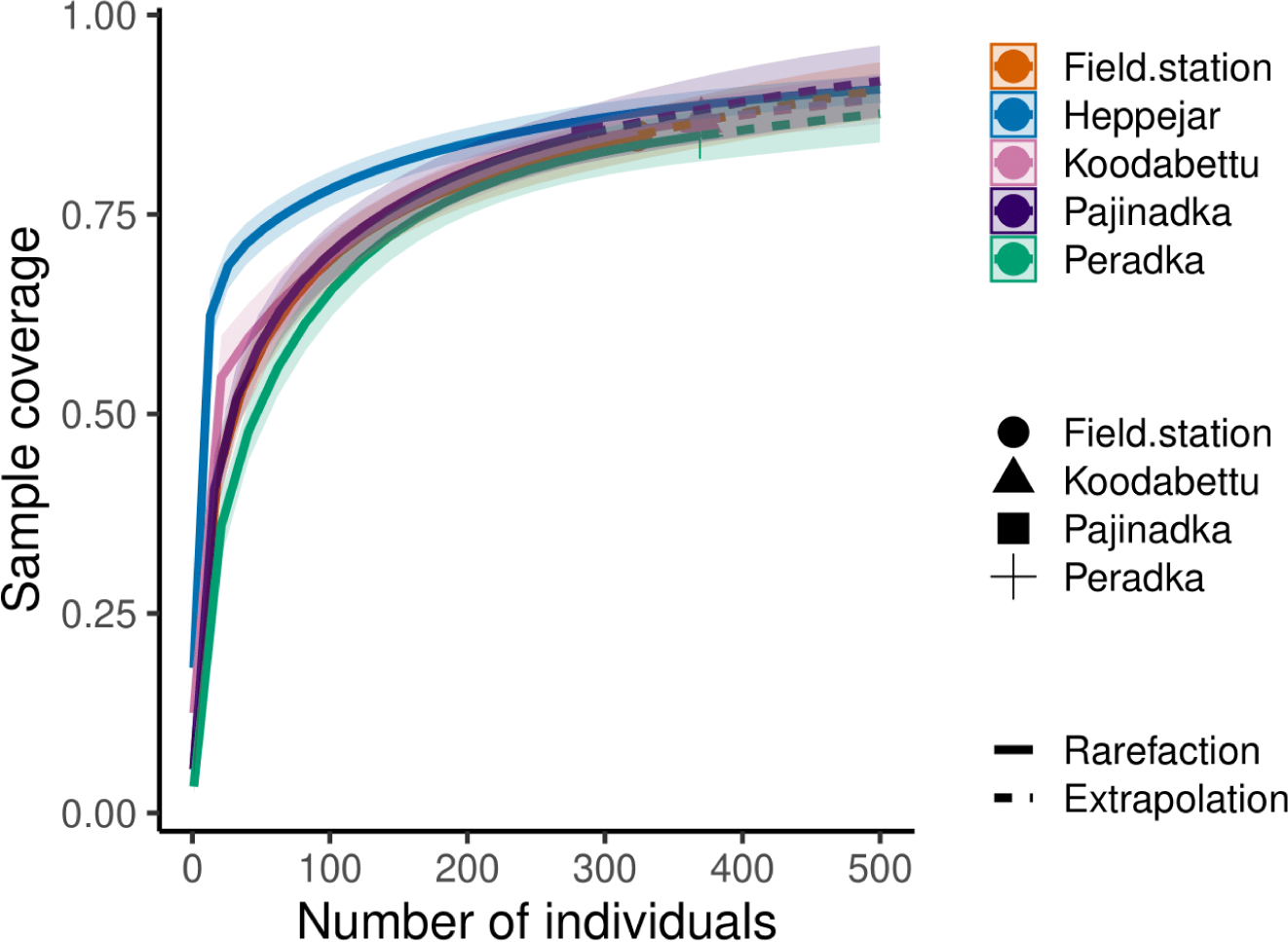
The graph shows an abundance-based rarefied and extrapolated species accumulation curve for the light trapping of moth species in the study area.

### Lepidoptera families consumed by M. spasma

Proportions of different families in the diet from the culled remains

In all the years, individuals of the family Sphingidae were found in proportionately higher numbers in the culled remains, followed by Erebidae, then Hepialidae (Fig. 4a). Geometridae species were found the least in Year 1; Limacodidae and Noctuidae in Year 2 and; Limacodidae and Zygaenidae in the last year.

**Figure 4:**
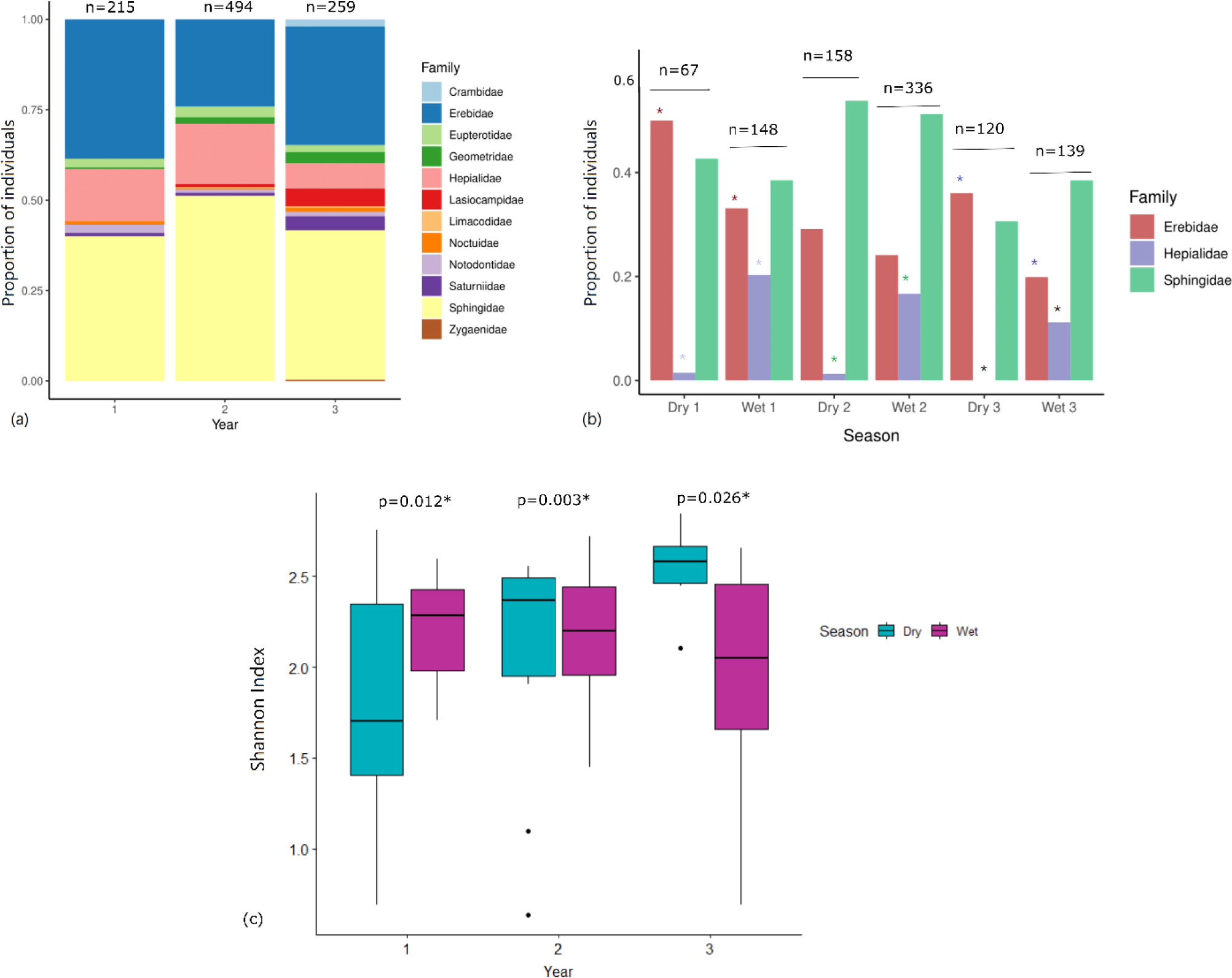
The proportion of individuals belonging to (a) different families of moths found in the culled remains across the years (b) Erebidae, Hepialidae, and Sphingidae families across seasons each year (* of similar colour denotes significant difference), (c) The diversity (Shannon Index) of moths found in the culled remains across the years in the Dry and Wet season (the plots show significant differences with Wilcoxon signed rank exact test)

The proportions of the most abundant families in the diet (Sphingidae, Erebidae and Hepialidae) in the diet varied significantly across dry and wet seasons in the sampling years. Erebids were eaten more in the dry season, in Year 1 and 2 and Hepialids were significantly eaten more in the wet season in all the years. Sphingids were found comparably in the diet throughout the seasons across years (Fig. 4b). We observed a significant difference in the overall diversity of moth species in the culled remains between the seasons each year (Fig. 4c).

Species belonging to the Saturniidae family had the largest wingspan (159.55±16.13 mm), forewing area (2856.45±667.8 mm^2^), hindwing area (2402.40±707.7 sq. mm), forewing length (88.60±10.27 mm), and hindwing length (66.02±11.49 mm). Hepialid moths showed the highest body length (54.55±5.45mm), Limacodids the highest Wing Loading (0.037±0.007), and Sphingidae moths showed the highest Forewing Aspect Ratio (4.00±0.02) and the lowest Maneuverability (0.52±0.005) (Fig. 5).

**Figure 5.**
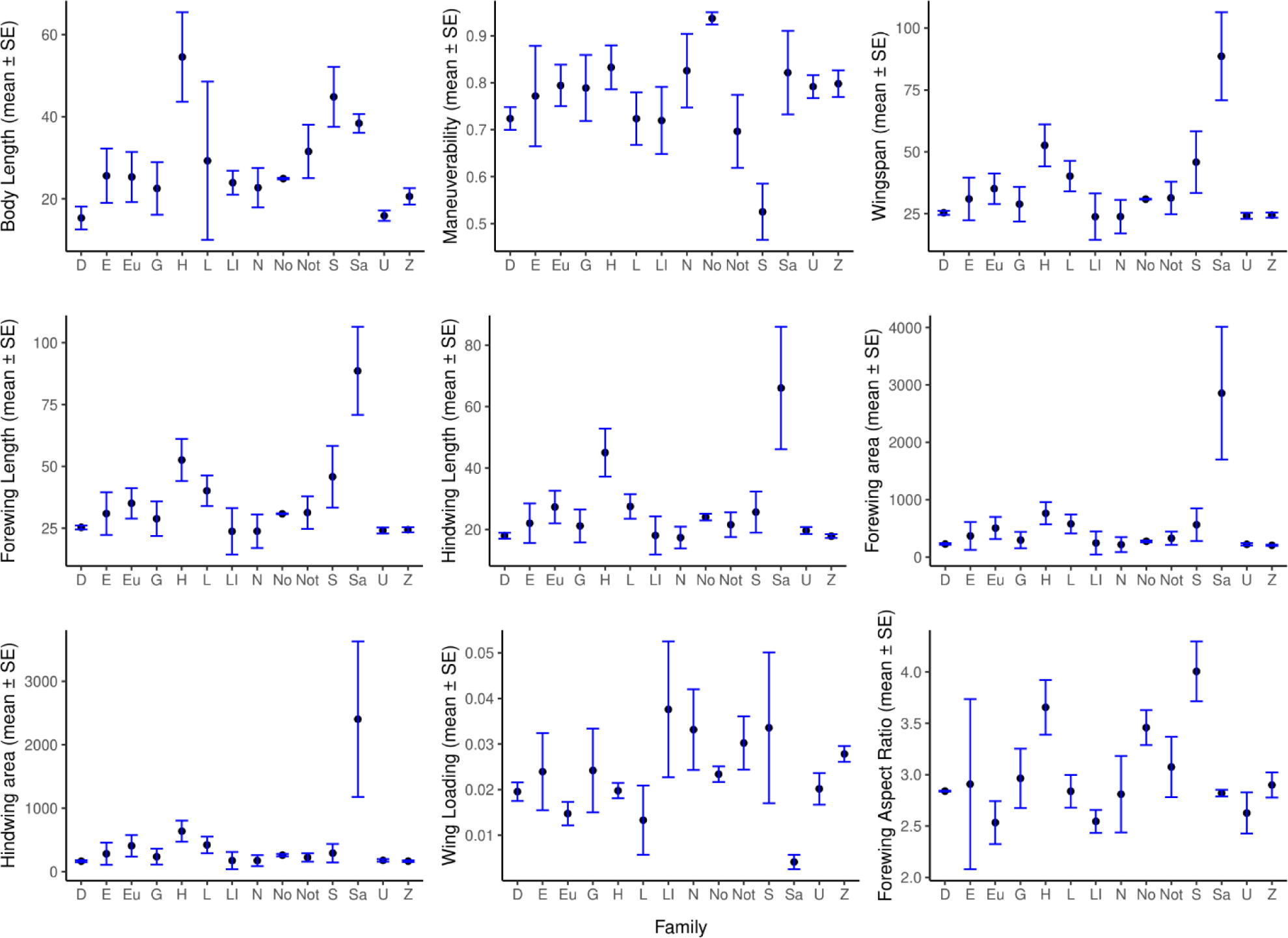
The 9 morphological trait values (in mm) (mean ± SE) measured across the families (wingspan >40mm) that were found in the diet. (D=Drepanidae, n=2; E=Erebidae, n=159; Eu=Eupterotidae, n=21; G=Geometridae, n=52; H=Hepialidae, n=4; L=Lasiocampidae, n=4; Li=Limacodidae, n=4; N=Noctuidae, n=12; No=Nolidae, n=2; Not=Notodontidae, n=22; S=Sphingidae, n=124; Sa=Saturniidae, n=3; U=Uraniidae, n=3; Z=Zygaenidae, n=5).

These traits were compared among the species present and absent in the diet (above ∼40 mm wingspan) from all the families. The species present in the diet were significantly larger, with higher Body Length, and other correlated traits like Wingspan, Forewing Area, Hindwing Area, Forewing Length and Hindwing length (Wilcoxon Rank sum test with continuity correction, p<0.01, for each of the traits). They also had higher Forewing Aspect Ratio (p<0.01), lower Wing Loading (p<0.01) and lower Maneuvrability (p <0.01) (Fig. 6).

**Figure 6.**
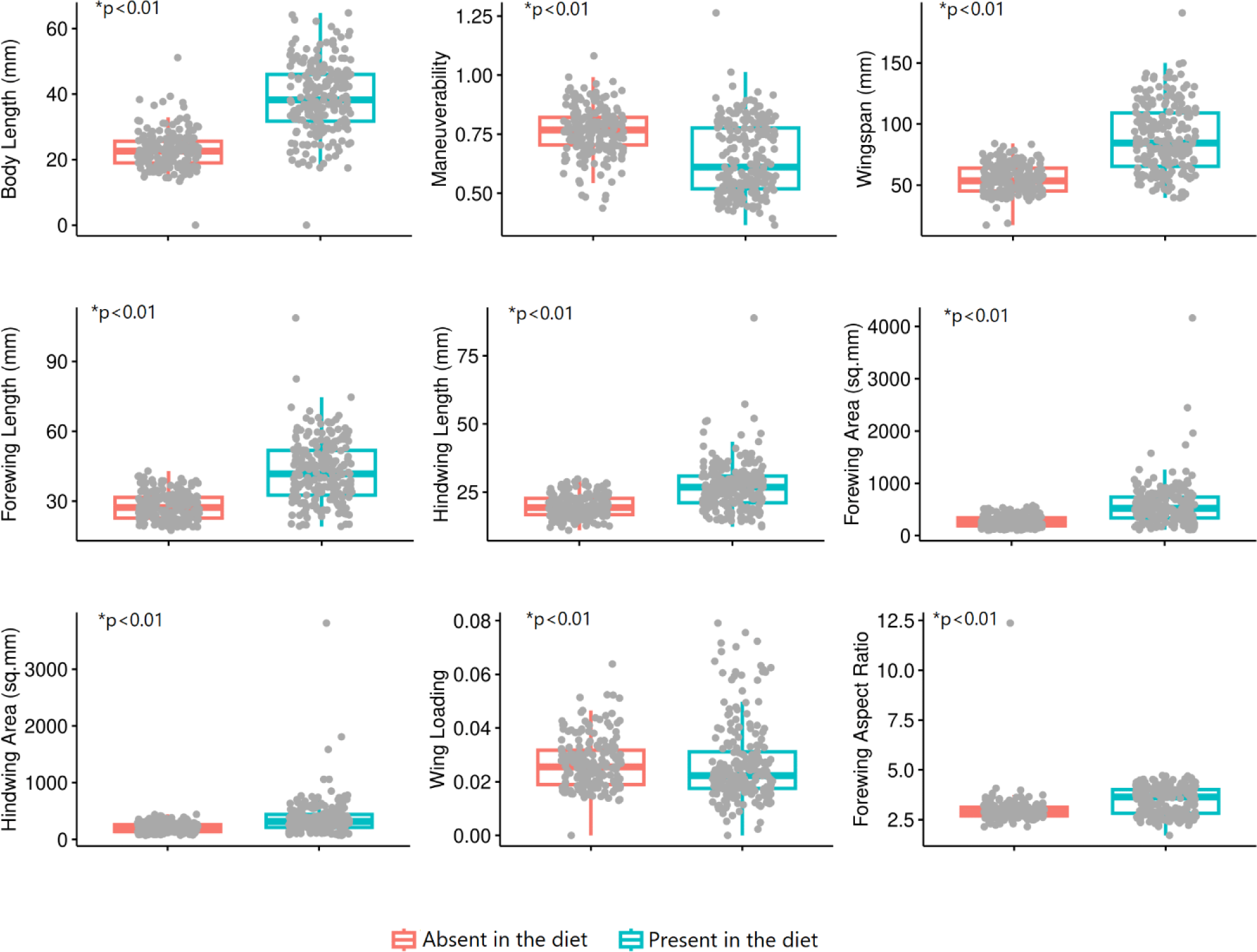
Comparison of the 9 morphological traits measured and pooled among the species that were present or absent in the diet of M. spasma (all the plots show significant differences with Wilcoxon Rank sum test).

There was a significant difference in the trait-space distances among the most abundant taxonomic groups found in the diet, i.e Sphinigidae, Erebidae and Hepialidae (PERMANOVA, df=1, R^2^=0.168, P<0.01) (Fig. 7a). Also, beta dispersion showed that the group dispersion, or variance, is significantly different from each other (p<0.01). When we looked into the traits of the species present or absent in the diet from these three taxonomic groups, we observed a significant difference in the trait-space distances (PERMANOVA, df=1, R^2^=0.274, P<0.01) (Fig. 7b) and the variance is significantly low (p=0.001) in the ‘absent’ group suggesting there is a specific range of traits that the bat is avoiding.

**Figure 7.**
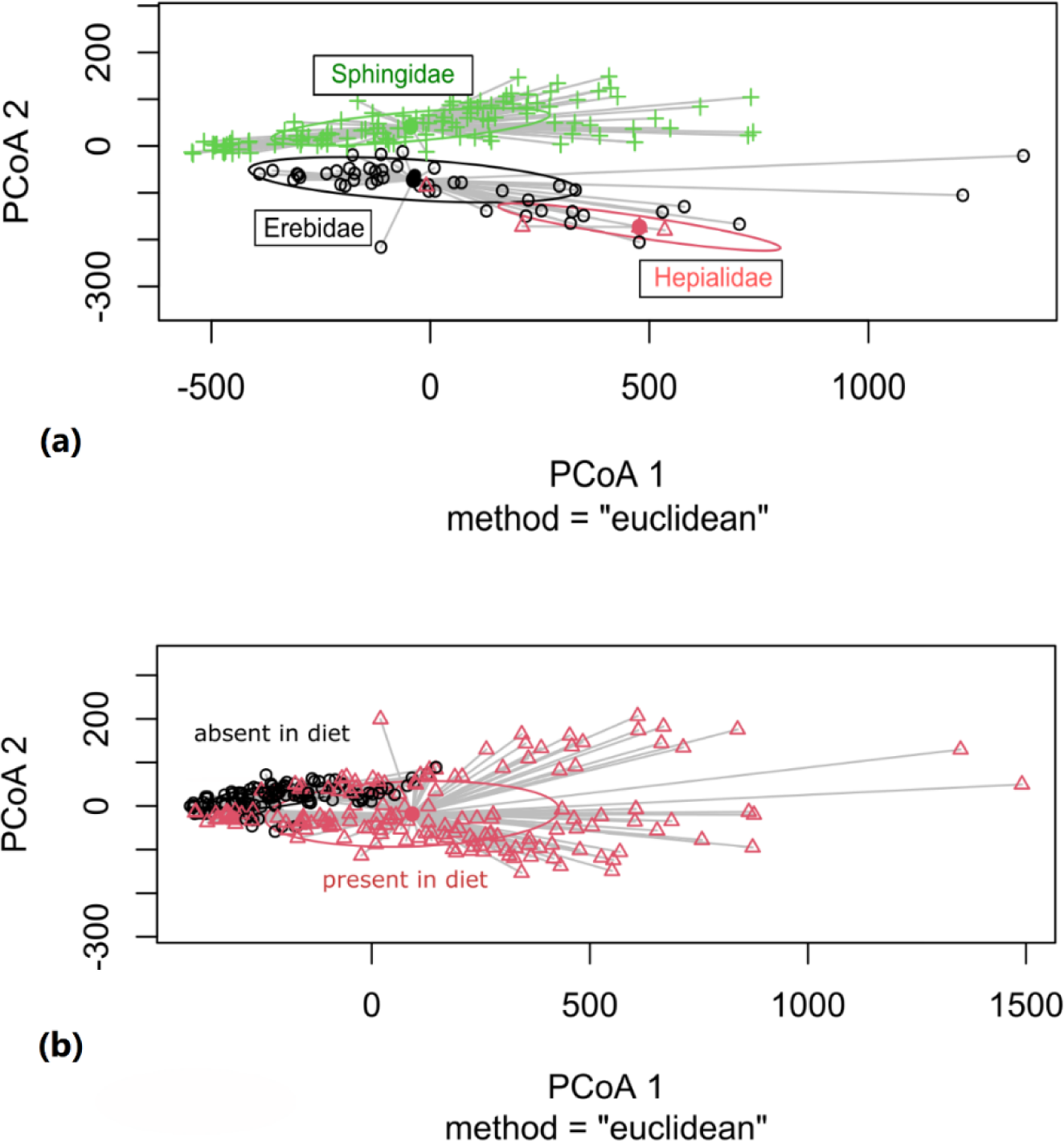
The Principal Coordinates Axes comparing the dispersion of traits a) the three most abundant families present in the diet (Erebidae, Sphingidae, and Hepialidae) and b) among species present and absent in the diet from these three subfamilies.

Body length (p<0.01, McFadden’s Pseudo R-square=0.45, Maneuverability (p<0.01, Mc Fadden’s Pseudo R-square=0.132), and Forewing Aspect Ratio (p<0.001, McFadden’s Pseudo R-square= 3.463261e-07), were found to be significant with Diet (present/absent) as a response variable in a Logistic regression model (model parameters in Supporting information Table 3). Predicted values from the Logistic Regression model showed that the higher the body size, Forewing Aspect Ratio, and lower the Maneuverability, there is increased probability of being present in the diet, wherein body size and maneuverability have the best fit (Fig. 8 a,b,c). Wing loading did not show any significant effect on the species being present or absent in the diet (Fig. 8d).

**Figure 8:**
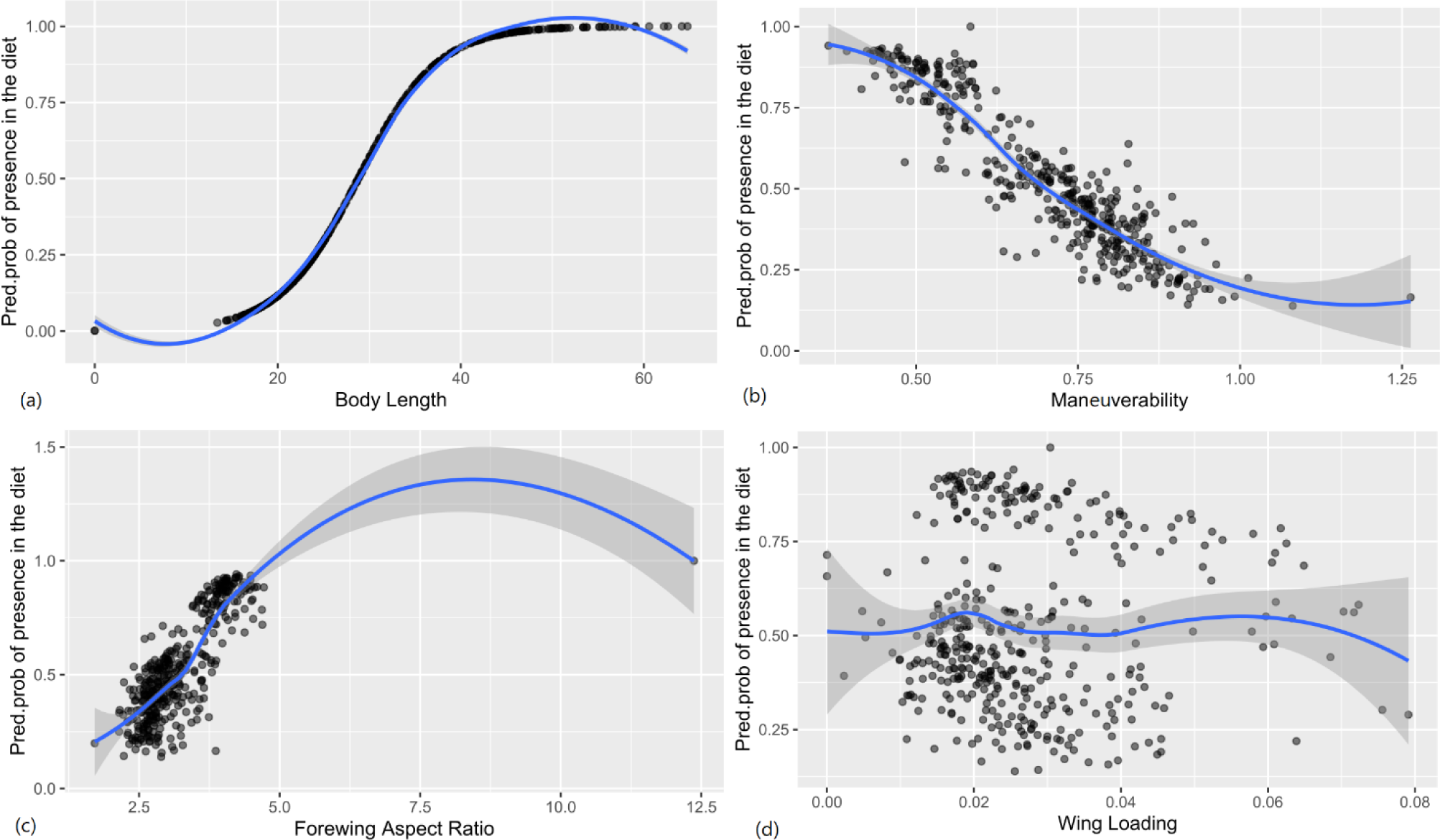
The logistic regression model plots show the predicted probabilities (with 95% CI) of the species to be present or absent in diet, with changing (a) Body Length, (b) Maneuverability, (c) Forewing Aspect Ratio, and (d) wing loading.

## Discussion

Our results show a seasonal variation in the overall moth species diversity in the diet of *M. spasma*, with Hepialid moths being preyed upon significantly more in the wet season in all the years. Our results also support the hypothesis that larger moths with weaker evasive capabilities are more vulnerable to predation by a sympatric bat predator. The study thus establishes a foundation for trait-based elucidation of the vulnerability of paleotropical moth communities to insectivorous bats.

### Seasonal variation in bat diet

Seasonal variation in diet is governed by food availability and how bats adapt to these changes (Bhartiy & Elangovan, 2021; Catto et al., 1995; Shiel et al., 1991) among others). Though Erebidae and Sphingidae were the most abundant moth families found overall in culled remains, we found seasonal variation in the composition of the diet of *M. spasma* across the years. This variation can be attributed to the availability of the moths of the family Hepialidae in the months of May and June every year (during the wet season) and also that they have the largest body size in our dataset. These moths emerge around the summer solstice and are known to be crepuscular or nocturnal (Andersson et al., 1998). Males are known to show lekking behaviour at dusk (for around 30 min) to avoid bird predators and late-emerging insectivorous bat species (Andersson et al., 1998; Rydell, 1998). Lekking behaviour in ‘acoustic’ moths is associated with reduced predation risk and increased mating success (Rydell, 1998), but the Hepialid moths are a primitive group of non-tympanate moths and do not produce ultrasound, mimetic coloration, or flight maneuvers to combat bat predation (Scoble, 1992), which places them at high risk of bat predation (Rydell, 1998). *M. spasma* is known to emerge at dusk, shortly after sunset (Balete, 2010; Prakash et al., 2021) and presumably hunts Hepialid moths due to their body size, and high detection during the months that they emerge, probably because of the male lekking behaviour. Amidst the global pandemic, due to a state-wide lockdown in the state of Karnataka, India, fieldwork could not be done in May and June 2021; hence we do not have light-trapping data from those months to account for the presence of these moths in the foraging area.

### Prey traits influencing vulnerability to predation

Insectivorous bats are known to detect and hunt prey in flight in a matter of seconds, during which the energy profit (Koselj et al., 2011), flight pattern, and ease of capture (Barclay & Brigham, 1994) of the prey would be prioritized over taxonomic predisposition (Arrizabalaga-Escudero et al., 2019). Although there was a shift in the local moth assemblage seasonally, our results show that prey body size is an important criterion for prey selection by *M. spasma*.

We did not find any Arctiinae moth body parts in the culled remains. These moths, commonly known as Tiger moths or Lichen moths are known to employ aposematic colouration and acoustic defence (warning, mimicry, or sonar-jamming) against bats (Miller & Surlykke, 2001; Ratcliffe et al., 2011; Ratcliffe & Fullard, 2005). Noctuid moths, also known for their anti-predatory flight maneuvers against bats (Roeder, 1962, 1964, 1967; Surlykke et al., 1999; ter Hofstede et al., 2013), scored pretty high on the maneuverability index (Fig. 4) and were less represented in the culled remains. Both these groups have >40mm wingspan and were abundant in our light traps.

Interestingly, moths of the Sphingidae family (commonly called hawkmoths) were recorded the most in the culled remains of *M. spasma*. Sphingid moths are known to have agile and maneuvering flight performance among flying insects (Greeter & Hedrick, 2016), with an increased ability to change the speed and direction of movement (Dudley, 2002). Such maneuverability may be beneficial for mate choice (Thornhill & Alcock, 2013) and defence. They show rapid turning-away flight when ‘visually’ startled to avoid ambush predators during flower visitation (Cheng et al., 2011; Wasserthal, 1993), but are not able to initiate escape flight as quickly as other ‘eared’ moths (that can detect bat echolocation), in response to bat echolocation calls (Morrill & Fullard, 1992). Here, using a morphological trait-based framework, we indeed find that Sphingids have the lowest hindwing-to-forewing area ratio, and highest forewing aspect ratio, which probably leads to less effective maneuvers against echolocating bats (correlation of form to function still needs detailed behavioural studies), and higher vulnerability to predation.

Also, certain species of hawkmoths (in two distantly related subtribes, the Choerocampina and the Acherontiina, (Göpfert et al., 2002) have non-tympanal hearing organs in the mouthparts that are capable of receiving ultrasound, but these groups were quite abundant in the culled remains in our results. Possibly, 1) Flying hawkmoths are detected by *M. spasma* from a longer distance due to their size (Surlykke et al., 1999), and audible wing beats (Balete, 2010), and 2) Hawkmoth ‘ears’ while flying or perching, are not able to detect the soft ‘low intensity’ echolocation calls of *M. spasma* (Fenton & Bell, 1981; Schmidt et al., 2000). Therefore, they are too late at initiating an escape flight.

Moths of the Erebidae family represented the second most abundant group in the diet of *M. spasma*. We found species from the genera *Erebus, Eudocima, Artena,* most commonly in the diet, which are the larger moths in the family, and possibly capable of anti-bat acoustic defence (Barber et al., 2022). Moths from the genus *Erebus* are not effectively attracted to light (Holloway, 2005) and hence were not attracted to our light traps, except in the months of August and September, (with few individuals) while moths of the genus *Artena* were quite abundant in our light traps. *Eudocima* moths are fruit-piercing moths and are generally found sucking on ripened citrus and other fruit crops (Reddy et al., 2007) during the evening, and could be easy targets for substrate-gleaning by bats. The study area is a human-dominated landscape with ample guava, cashewnut, *Areca*, and other fruiting trees, which are common hosts of the fruit piercers. The high occurrence of these groups in the culled remains could be due to their abundance in the foraging range of the bats and easy detectability by *M. spasma* (large size with audible wingbeats, (Balete, 2010). Not much has been established in the literature to clarify the acoustic or behavioural defences against bats for these groups of moths.

### Prey selection by the bat predator

Our study showed some degree of selective hunting of moth species, previously unknown for a megadermatid bat, which is not a moth specialist. *M. spasma* and related bats with similar foraging strategies rely more on prey-generated sound for prey detection at long range (Balete, 2010; Denzinger & Schnitzler, 2013; Raghuram et al., 2015; Schnitzler et al., 2003), but can also hunt through echolocation similar to its closest relative *Lyroderma lyra* (Marimuthu et al., 1995; Ratcliffe et al., 2005; Schmidt et al., 2000). Given the broad range in diet (Raghuram et al., 2015), this bat most likely adopts a flexible strategy of hawking and gleaning (Gordon et al., 2019). In our study, it was, however, not possible to establish if the moths found in the diet were gleaned from a surface or caught in the air.

Even though the diet might be largely conserved in sympatric predators, each species might segregate temporally, spatially, or through different hunting strategies (Divoll et al., 2022). How is the trophic niche of moths shaped in this landscape, in adaptation to the range of hunting strategies of different species of their most formidable predator? Following (Arrizabalaga-Escudero et al., 2019); using both prey and predator traits, we aim in future studies to generate a deeper understanding of prey selection in this biodiverse landscape. In the face of the alarming decline of insect populations (Wagner et al., 2021), we could potentially identify predators that are affected by the loss of key prey traits when there is a decline in prey populations.

## Supplementary material

**Table 1:**
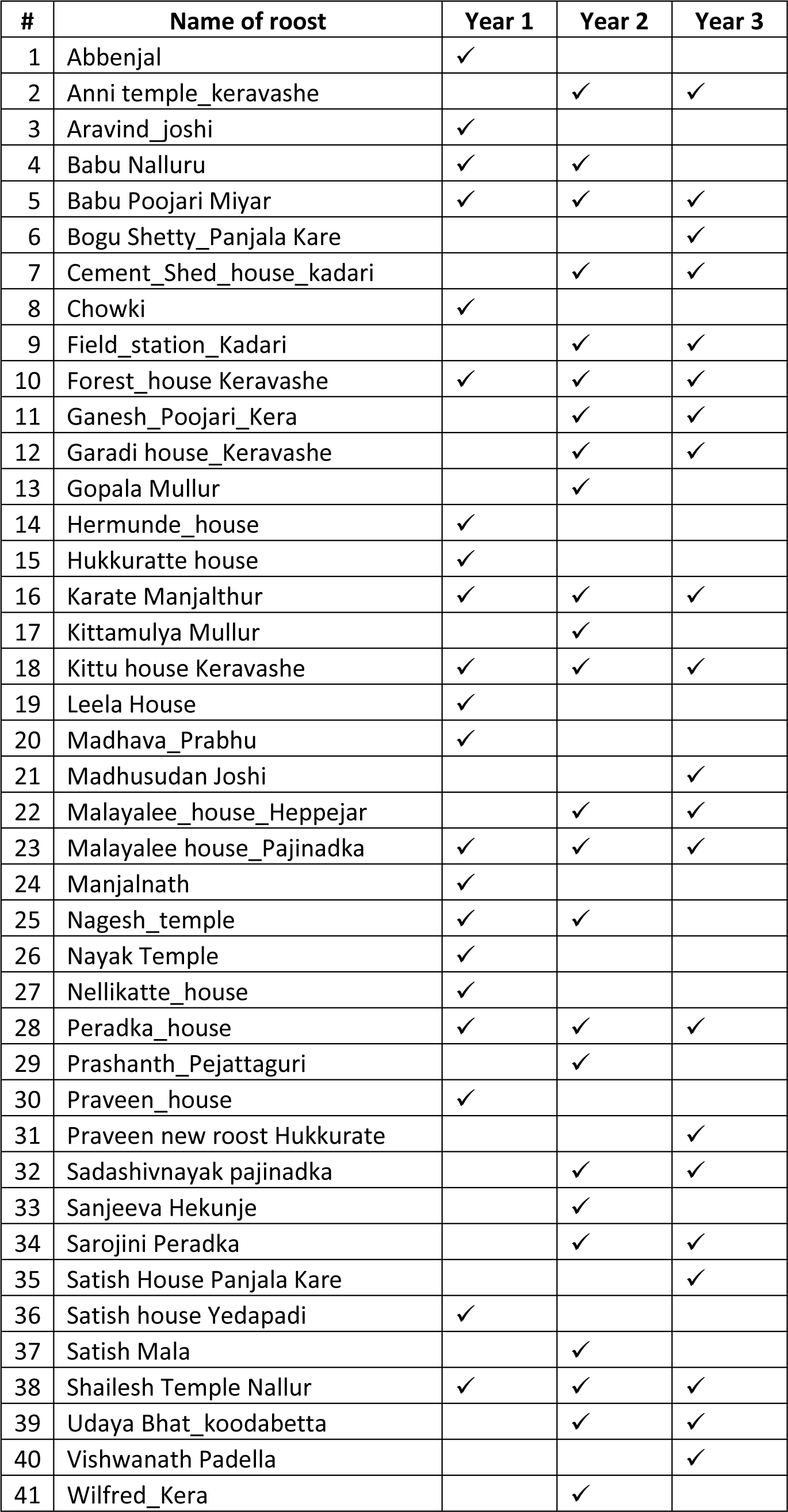
The bat roosts sampled across the years.

| # | Name of roost | Year 1 | Year 2 | Year 3 |
| --- | --- | --- | --- | --- |
| 1 | Abbenjal | ✓ |  |  |
| 2 | Anni temple_keravashe |  | ✓ | ✓ |
| 3 | Aravind_joshi | ✓ |  |  |
| 4 | Babu Nalluru | ✓ | ✓ |  |
| 5 | Babu Poojari Miyar | ✓ | ✓ | ✓ |
| 6 | Bogu Shetty_Panjala Kare |  |  | ✓ |
| 7 | Cement_Shed_house_kadari |  | ✓ | ✓ |
| 8 | Chowki | ✓ |  |  |
| 9 | Field_station_Kadari |  | ✓ | ✓ |
| 10 | Forest_house Keravashe | ✓ | ✓ | ✓ |
| 11 | Ganesh_Poojari_Kera |  | ✓ | ✓ |
| 12 | Garadi house_Keravashe |  | ✓ | ✓ |
| 13 | Gopala Mullur |  | ✓ |  |
| 14 | Hermunde_house | ✓ |  |  |
| 15 | Hukkuratte house | ✓ |  |  |
| 16 | Karate Manjalthur | ✓ | ✓ | ✓ |
| 17 | Kittamulya Mullur |  | ✓ |  |
| 18 | Kittu house Keravashe | ✓ | ✓ | ✓ |
| 19 | Leela House | ✓ |  |  |
| 20 | Madhava_Prabhu | ✓ |  |  |
| 21 | Madhusudan Joshi |  |  | ✓ |
| 22 | Malayalee_house_Heppejar |  | ✓ | ✓ |
| 23 | Malayalee house_Pajinadka | ✓ | ✓ | ✓ |
| 24 | Manjalnath | ✓ |  |  |
| 25 | Nagesh_temple | ✓ | ✓ |  |
| 26 | Nayak Temple | ✓ |  |  |
| 27 | Nellikatte_house | ✓ |  |  |
| 28 | Peradka_house | ✓ | ✓ | ✓ |
| 29 | Prashanth_Pejattaguri |  | ✓ |  |
| 30 | Praveen_house | ✓ |  |  |
| 31 | Praveen new roost Hukkurate |  |  | ✓ |
| 32 | Sadashivnayak pajinadka |  | ✓ | ✓ |
| 33 | Sanjeeva Hekunje |  | ✓ |  |
| 34 | Sarojini Peradka |  | ✓ | ✓ |
| 35 | Satish House Panjala Kare |  |  | ✓ |
| 36 | Satish house Yedapadi | ✓ |  |  |
| 37 | Satish Mala |  | ✓ |  |
| 38 | Shailesh Temple Nallur | ✓ | ✓ | ✓ |
| 39 | Udaya Bhat_koodabetta |  | ✓ | ✓ |
| 40 | Vishwanath Padella |  |  | ✓ |
| 41 | Wilfred_Kera |  | ✓ |  |

**Figure S1:**
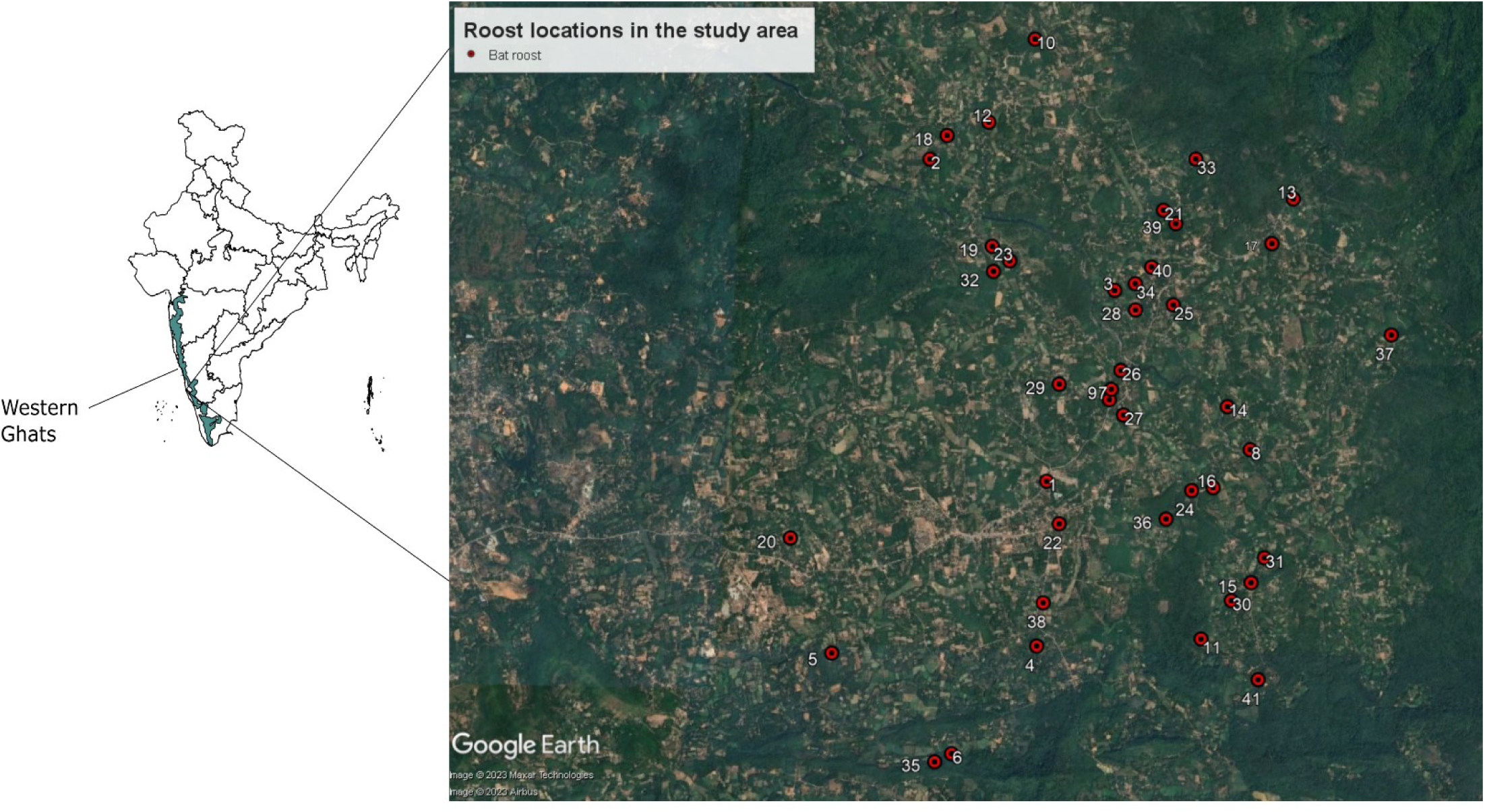
Spatial roost locations for culled remains collection in the study area. The numbers correspond to the roost numbers in the Table 1.

**Table 2:** Details of the light traps.

| Sites | Total no. of nights sampled (sampling months) | Plots | Plot type | Total no. of species recorded |
| --- | --- | --- | --- | --- |
| Field station | 14 (Jan-April; Aug-Oct) | P1 | Forest | 94 |
|  |  | P2 | Plantation |  |
|  |  | P3 | Forest |  |
|  |  | P4 | Forest |  |
| Peradka | 14 (Jan-April; Aug-Oct) | P1 | Forest | 111 |
|  |  | P2 | Forest |  |
|  |  | P3 | Forest |  |
|  |  | P4 | Forest |  |
| Pajinadka | 14 (Jan-April; Aug-Oct) | P1 | Forest | 85 |
|  |  | P2 | Plantation |  |
|  |  | P3 | Forest |  |
|  |  | P4 | Forest |  |
| Koodabettu | 14 (Jan-April; Aug-Oct) | P1 | Plantation | 94 |
|  |  | P2 | Forest |  |
|  |  | P3 | Forest |  |
|  |  | P4 | Forest |  |
| Heppejar | 14 (Jan-April; Aug-Oct) | P1 | Plantation | 100 |
|  |  | P2 | Forest |  |
|  |  | P3 | Forest |  |
|  |  | P4 | Forest |  |

**Figure S2:**
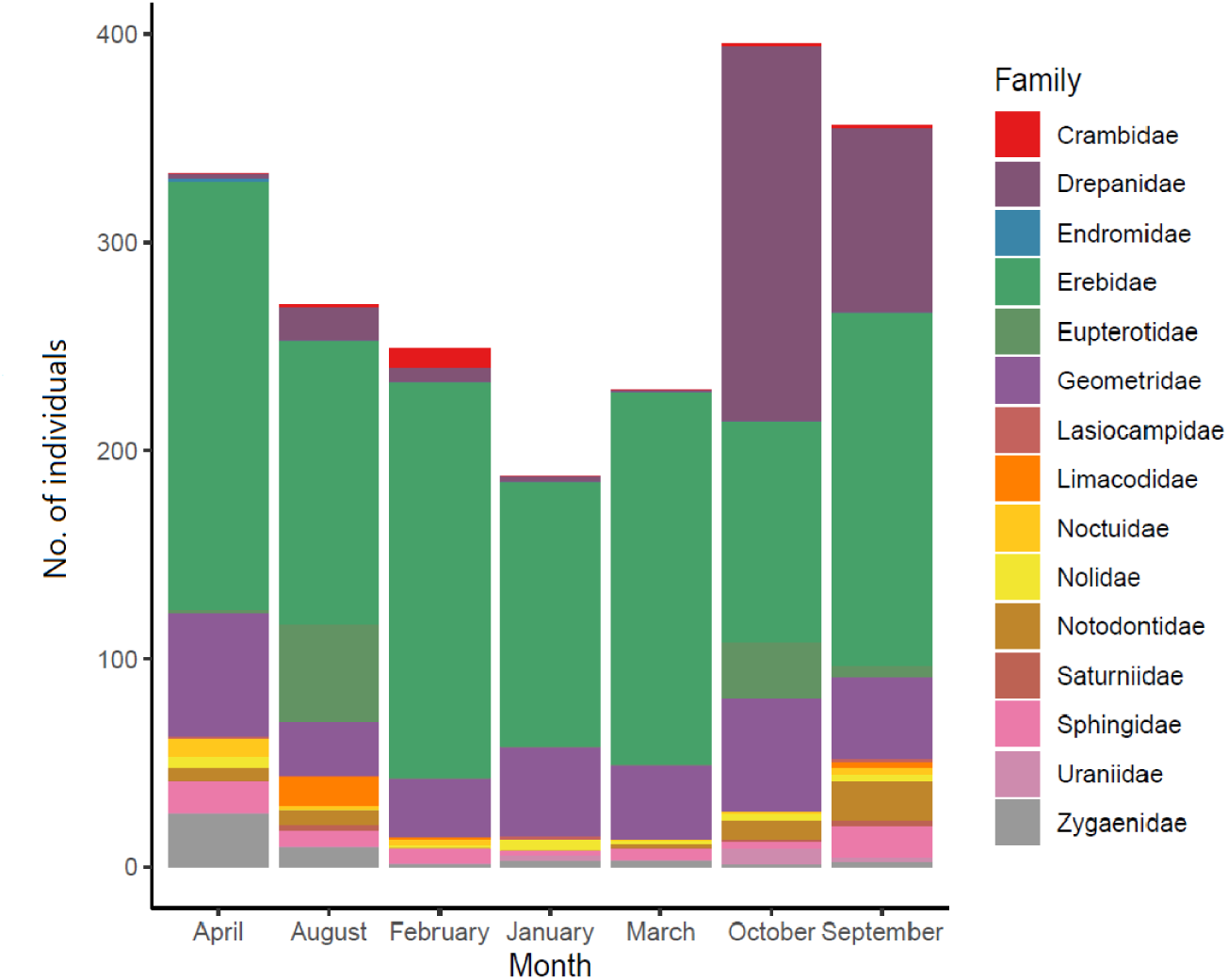
The number of individuals from each family (>40mm wingspan) counted at the light traps across 7 months in 2021.

**Figure S3:**
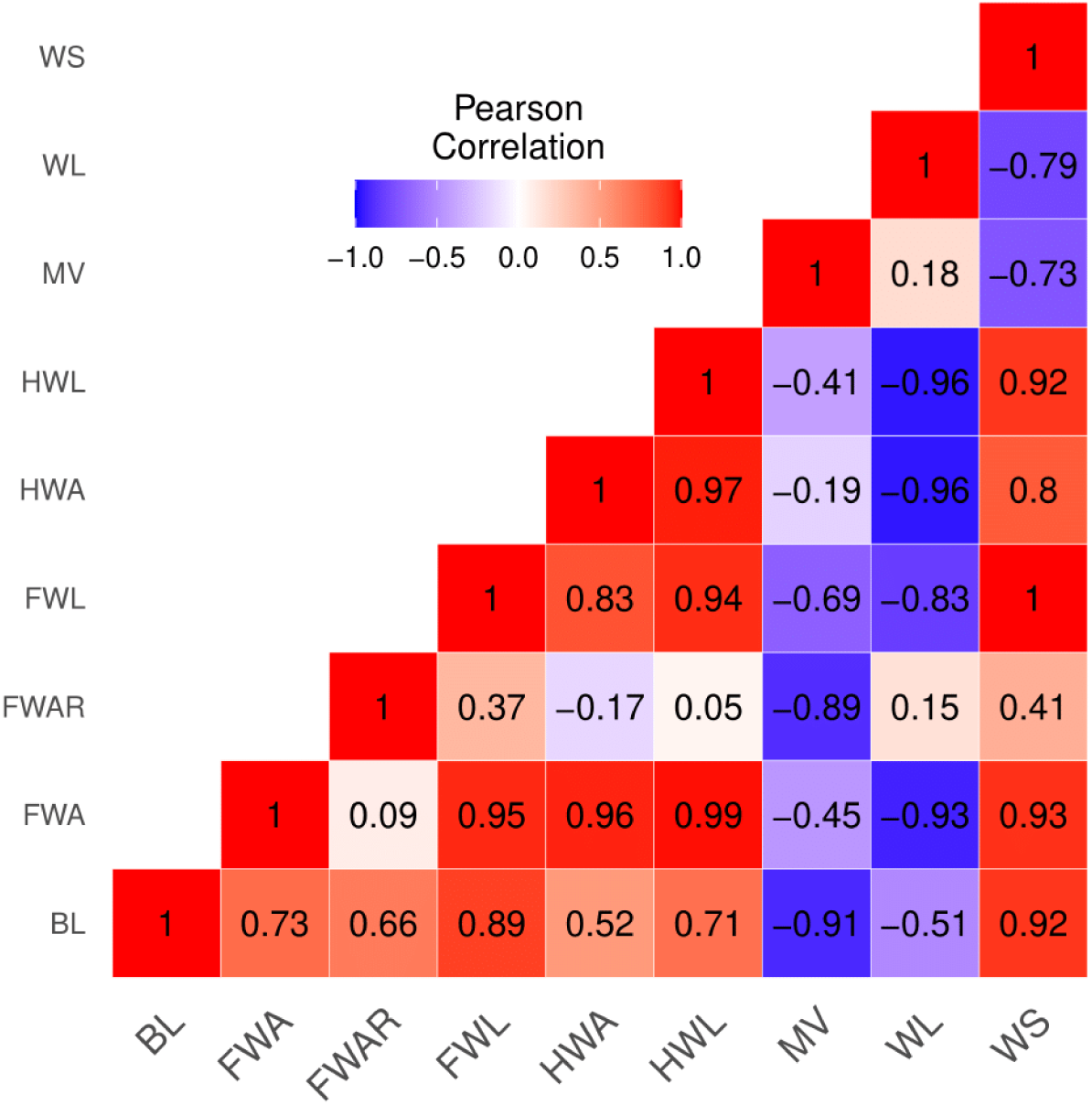
Correlation matrix for the morphological traits measured from the moth specimens (Forewing length (FWL); hindwing length (HWL); Body length (BL) (a proxy for body size); Wingspan; Forewing area (FWA); Hindwing area (HWA); Maneuverability=HWA/FWA; Wingloading=BL/2(FWA+HWA) and Forewing aspect ratio = (2*FWL)/FWA (as a proxy for wing shape)

**Table 3:** Model parameters for the Logistic Regression models (binomial function) for each of the traits.

| Trait | Model | Estimate | std. error | Z-value | p-value |
| --- | --- | --- | --- | --- | --- |
| Body Length | glm(formula = Diet ~ BL, family = binomial, data = Trait_family) | 0.2309 | 0.0221 | 10.44 | 0.001 |
| Hindwing Area/Forewing Area=Maneuverability | glm(formula = Diet ~ MV, family = binomial, data = Trait_family) | -6.7912 | 0.8689 | -7.816 | 0.001 |
| (2*Forewing Length)/Forewing Area =Forewing Aspect Ratio | glm(formula = Diet ~ FWAR, family = binomial, data = Trait_family) | 1.3809 | 0.1924 | 7.179 | 0.001 |
| Body Length/2(Forewing Area+Hindwing Area)=Wing Loading | glm(formula = Diet ~ WL, family = binomial, data = Trait_family) | 0.11 | 7.7737 | 0.014 | 0.98 |

## References

Almenar, D., Aihartza, J., Goiti, U., Salsamendi, E., & Garin, I. (2008). Diet and prey selection in the trawling long-fingered bat. Journal of Zoology, 274(4), 340–348. https://doi.org/10.1111/j.1469-7998.2007.00390.x

Almenar, D., Aihartza, J., Goiti, U., Salsamendi, E., & Garin, I. (2013). Hierarchical patch choice by an insectivorous bat through prey availability components. Behavioral Ecology and Sociobiology, 67(2), 311–320. https://doi.org/10.1007/s00265-012-1451-z

Andersson, S., Rydell, J., & Svensson, M. G. E. (1998). Light, predation and the lekking behaviour of the ghost swift *Hepialus humuli* (L.) (Lepidoptera, Hepialidae). Proceedings of the Royal Society of London. Series B: Biological Sciences, 265(1403), 1345–1351. https://doi.org/10.1098/rspb.1998.0440

Arrizabalaga-Escudero, A., Clare, E. L., Salsamendi, E., Alberdi, A., Garin, I., Aihartza, J., & Goiti, U. (2018). Assessing niche partitioning of co-occurring sibling bat species by DNA metabarcoding. Molecular Ecology, 27(5), 1273–1283. https://doi.org/10.1111/mec.14508

Arrizabalaga-Escudero, A., Merckx, T., García-Baquero, G., Wahlberg, N., Aizpurua, O., Garin, I., Goiti, U., & Aihartza, J. (2019). Trait-based functional dietary analysis provides a better insight into the foraging ecology of bats. Journal of Animal Ecology, 88(10), 1587–1600. https://doi.org/10.1111/1365-2656.13055

Balete, D. (2010). Food and roosting habits of the lesser false vampire bat, *Megaderma spasma* (Chiroptera: Megadermatidae) in a Philippine lowland forest. Asia Life Sciences, 4, 111–129.

Barber, J. R., Plotkin, D., Rubin, J. J., Homziak, N. T., Leavell, B. C., Houlihan, P. R., Miner, K. A., Breinholt, J. W., Quirk-Royal, B., Padrón, P. S., Nunez, M., & Kawahara, A. Y. (2022). Anti-bat ultrasound production in moths is globally and phylogenetically widespread. Proceedings of the National Academy of Sciences, 119(25), e2117485119. https://doi.org/10.1073/pnas.2117485119

Barclay, R. M. R., & Brigham, R. M. (1994). Constraints on optimal foraging: A field test of prey discrimination by echolocating insectivorous bats. Animal Behaviour, 48(5), 1013–1021. https://doi.org/10.1006/anbe.1994.1334

Basset, Y., Cizek, L., Cuénoud, P., Didham, R. K., Guilhaumon, F., Missa, O., Novotny, V., Ødegaard, F., Roslin, T., Schmidl, J., Tishechkin, A. K., Winchester, N. N., Roubik, D. W., Aberlenc, H.-P., Bail, J., Barrios, H., Bridle, J. R., Castaño-Meneses, G., Corbara, B., … Leponce, M. (2012). Arthropod Diversity in a Tropical Forest. Science, 338(6113), 1481–1484. https://doi.org/10.1126/science.1226727

Bhartiy, S. K., & Elangovan, V. (2021). Seasonal prey availability and diet composition of Lesser Asiatic Yellow House Bat *Scotophilus kuhlii* Leach, 1821. Journal of Threatened Taxa, 13(8), 19002–19010. https://doi.org/10.11609/jott.6296.13.8.19002-19010

Brehm, G. (2017). A new LED lamp for the collection of nocturnal Lepidoptera and a spectral comparison of light-trapping lamps. Nota Lepidopterologica, 40(1), 87–108. https://doi.org/10.3897/nl.40.11887

Brose, U. (2010). Body-mass constraints on foraging behaviour determine population and food-web dynamics. Functional Ecology, 24(1), 28–34. https://doi.org/10.1111/j.1365-2435.2009.01618.x

Brosset, A. (1962). The bats of central and western India. Journal of Bombay Natural History Society, 59: 1–78.

Burles, D. W., Brigham, R. M., Ring, R. A., & Reimchen, T. E. (2008). Diet of two insectivorous bats, *Myotis lucifugus* and *Myotis keenii*, in relation to arthropod abundance in a temperate Pacific Northwest rainforest environment. Canadian Journal of Zoology, 86(12), 1367–1375. https://doi.org/10.1139/Z08-125

Catania, K. C., & Remple, F. E. (2005). Asymptotic prey profitability drives star-nosed moles to the foraging speed limit. Nature, 433(7025), 519–522. https://doi.org/10.1038/nature03250

Catto, C. M. C., Racey, P. A., & Stephenson, P. J. (1995). Activity patterns of the serotine bat (*Eptesicus serotinus*) at a roost in southern England. Journal of Zoology, 235(4), 635–644. https://doi.org/10.1111/j.1469-7998.1995.tb01774.x

Cheng, B., Deng, X., & Hedrick, T. L. (2011). The mechanics and control of pitching manoeuvres in a freely flying hawkmoth (*Manduca sexta*). Journal of Experimental Biology, 214(24), 4092–4106. https://doi.org/10.1242/jeb.062760

Davison, G. W. H., & Zubaid, A. (1992). Food habits of the Lesser false vampire, *Megaderma spasma*, from Kuala Lompat, Peninsular Malaysia. Zeitschrift für Säugetierkunde, 57(5), 310–312.

Denzinger, A., & Schnitzler, H.U. (2013). Bat guilds, a concept to classify the highly diverse foraging and echolocation behaviors of microchiropteran bats. Frontiers in Physiology, 4, 1–15. https://www.frontiersin.org/articles/10.3389/fphys.2013.00164

Divoll, T. J., Brown, V. A., McCracken, G. F., & O’Keefe, J. M. (2022). Prey size is more representative than prey taxa when measuring dietary overlap in sympatric forest bats. Environmental DNA, edn3.354. https://doi.org/10.1002/edn3.354

Dudley, R. (2002). Mechanisms and Implications of Animal Flight Maneuverability. Integrative and Comparative Biology, 42(1), 135–140. https://doi.org/10.1093/icb/42.1.135

Fenton, M. B., & Bell, G. P. (1981). Recognition of Species of Insectivorous Bats by Their Echolocation Calls. Journal of Mammalogy, 62(2), 233–243. https://doi.org/10.2307/1380701

Fenton, M. B., Swanepoel, C. M., Brigham, R. M., Cebek, J., & Hickey, M. B. C. (1990). Foraging Behavior and Prey Selection by Large Slit-Faced Bats (*Nycteris grandis*; Chiroptera: Nycteridae). Biotropica, 22(1), 2–8. https://doi.org/10.2307/2388713

Ferry-Graham, L. A. (2002). Using Functional Morphology to Examine the Ecology and Evolution of Specialization. Integrative and Comparative Biology, 42(2), 265–277. https://doi.org/10.1093/icb/42.2.265

Göpfert, M. C., Surlykke, A., & Wasserthal, L. T. (2002). Tympanal and atympanal “mouth-ears” in hawkmoths (Sphingidae). Proceedings of the Royal Society B: Biological Sciences, 269(1486), 89–95. https://doi.org/10.1098/rspb.2001.1646

Gordon, R., Ivens, S., Ammerman, L. K., Fenton, M. B., Littlefair, J. E., Ratcliffe, J. M., & Clare, E. L. (2019). Molecular diet analysis finds an insectivorous desert bat community dominated by resource sharing despite diverse echolocation and foraging strategies. Ecology and Evolution, 9(6), 3117–3129. https://doi.org/10.1002/ece3.4896

Greeter, J. S. M., & Hedrick, T. L. (2016). Direct lateral maneuvers in hawkmoths. Biology Open, 5(1), 72–82. https://doi.org/10.1242/bio.012922

Holloway, J.D. (1997). The moths of Borneo (vol.15-16): Family Noctuidae, subfamily Catocalinae. Malayan Nature Journal, 51, 1–242.

Howland, H. C. (1974). Optimal strategies for predator avoidance: The relative importance of speed and manoeuvrability. Journal of Theoretical Biology, 47(2), 333–350. https://doi.org/10.1016/0022-5193(74)90202-1

Hsieh, T. C., Ma, K. H., & Chao, A. (2016). iNEXT: An R package for rarefaction and extrapolation of species diversity (Hill numbers). Methods in Ecology and Evolution, 7(12), 1451–1456. https://doi.org/10.1111/2041-210X.12613

Hügel, T., & Goerlitz, H. R. (2019). Species-specific strategies increase unpredictability of escape flight in eared moths. Functional Ecology, 33(9), 1674–1683. https://doi.org/10.1111/1365-2435.13383

Jantzen, B., & Eisner, T. (2008). Hindwings are unnecessary for flight but essential for execution of normal evasive flight in Lepidoptera. Proceedings of the National Academy of Sciences, 105(43), 16636–16640. https://doi.org/10.1073/pnas.0807223105

Jones, G. (1990). Prey Selection by the Greater Horseshoe Bat (*Rhinolophus ferrumequinum*): Optimal Foraging by Echolocation? Journal of Animal Ecology, 59(2), 587–602. https://doi.org/10.2307/4882

Klecka, J., & Boukal, D. S. (2013). Foraging and vulnerability traits modify predator–prey body mass allometry: Freshwater macroinvertebrates as a case study. Journal of Animal Ecology, 82(5), 1031–1041. https://doi.org/10.1111/1365-2656.12078

Koselj, K., Schnitzler, H. U., & Siemers, B. M. (2011). Horseshoe bats make adaptive prey-selection decisions, informed by echo cues. Proceedings of the Royal Society B: Biological Sciences, 278(1721), 3034–3041. https://doi.org/10.1098/rspb.2010.2793

Marimuthu, G., Habersetzer, J., & Leippert, D. (1995). Active Acoustic Gleaning from the Water Surface by the Indian False Vampire Bat, *Megaderma lyra*. Ethology, 99(1–2), 61–74. https://doi.org/10.1111/j.1439-0310.1995.tb01089.x

Mata, V. A., Amorim, F., Corley, M. F. V., McCracken, G. F., Rebelo, H., & Beja, P. (2016). Female dietary bias towards large migratory moths in the European free-tailed bat (*Tadarida teniotis*). Biology Letters, 12(3), 20150988. https://doi.org/10.1098/rsbl.2015.0988

Miller, L. A., & Surlykke, A. (2001). How Some Insects Detect and Avoid Being Eaten by Bats: Tactics and Countertactics of Prey and Predator. BioScience, 51(7), 570–581. https://doi.org/10.1641/0006-3568(2001)051[0570:HSIDAA]2.0.CO;2

Minnaar, C., Boyles, J. G., Minnaar, I. A., Sole, C. L., & McKechnie, A. E. (2015). Stacking the odds: Light pollution may shift the balance in an ancient predator–prey arms race. Journal of Applied Ecology, 52(2), 522–531. https://doi.org/10.1111/1365-2664.12381

Moore, T. Y., & Biewener, A. A. (2015). Outrun or Outmaneuver: Predator–Prey Interactions as a Model System for Integrating Biomechanical Studies in a Broader Ecological and Evolutionary Context. Integrative and Comparative Biology, 55(6), 1188–1197. https://doi.org/10.1093/icb/icv074

Morrill, S. B., & Fullard, J. H. (1992). Auditory influences on the flight behaviour of moths in a Nearctic site. I. Flight tendency. Canadian Journal of Zoology, 70(6), 1097–1101. https://doi.org/10.1139/z92-153

Nagaraja, B.C., Somashekar, R. K., & Raj, B. (2005). Tree species diversity and composition in logged and unlogged rainforest of Kudremukh National Park, South India. Journal of Environmental Biology / Academy of Environmental Biology, India, 26, 627–634.

Nakazawa, T. (2017). Individual interaction data are required in community ecology: A conceptual review of the predator–prey mass ratio and more. Ecological Research, 32(1), 5–12. https://doi.org/10.1007/s11284-016-1408-1

Oksanen, J., Blanchet, F.G., Kindt, R., Legendre, P., Minchin, P.R., O’hara, R.B., Simpson, G.L., Solymos, P., Stevens, M.H.H., Wagner, H. and Oksanen, M.J. (2013). Package ‘vegan’. Community ecology package, version, 2(9), 1–295.

Ortiz, E., & Arim, M. (2016). Hypotheses and trends on how body size affects trophic interactions in a guild of South American killifishes. Austral Ecology, 41(8), 976–982. https://doi.org/10.1111/aec.12389

Pascal, J. P., Ramesh, B. R., & Franceschi, D. D. (2004). Wet evergreen forest types of the southern western ghats, India. Tropical Ecology 45(2),281–292

Prakash, H. (2020). *Foraging decisions of the Lesser False Vampire Bat, Megaderma spasma in a heterogenous landscape* (PhD Thesis). Indian Institute of Science

Prakash, H., Saha, K., Sahu, S., & Balakrishnan, R. (2021). Ecological drivers of selection for remnant forest habitats by an insectivorous bat in a tropical, human-modified landscape. Forest Ecology and Management, 496, 119451. https://doi.org/10.1016/j.foreco.2021.119451

Raghuram, H., Deb, R., Nandi, D., & Balakrishnan, R. (2015). Silent katydid females are at higher risk of bat predation than acoustically signalling katydid males. Proceedings of the Royal Society B: Biological Sciences, 282(1798), 20142319. https://doi.org/10.1098/rspb.2014.2319

Ratcliffe, J. M., & Fullard, J. H. (2005). The adaptive function of tiger moth clicks against echolocating bats: An experimental and synthetic approach. Journal of Experimental Biology, 208(24), 4689–4698. https://doi.org/10.1242/jeb.01927

Ratcliffe, J. M., Fullard, J. H., Arthur, B. J., & Hoy, R. R. (2011). Adaptive auditory risk assessment in the dogbane tiger moth when pursued by bats. Proceedings of the Royal Society B: Biological Sciences, 278(1704), 364–370. https://doi.org/10.1098/rspb.2010.1488

Ratcliffe, J. M., Raghuram, H., Marimuthu, G., Fullard, J. H., & Fenton, M. B. (2005). Hunting in unfamiliar space: Echolocation in the Indian false vampire bat, *Megaderma lyra*, when gleaning prey. Behavioral Ecology and Sociobiology, 58(2), 157–164. https://doi.org/10.1007/s00265-005-0912-z

Reddy, G. V. P., Cruz, Z. T., & Muniappan, R. (2007). Attraction of fruit-piercing moth *Eudocima phalonia* (Lepidoptera: Noctuidae) to different fruit baits. Crop Protection, 26(4), 664–667. https://doi.org/10.1016/j.cropro.2006.06.004

Roeder, K. D. (1962). The behaviour of free flying moths in the presence of artificial ultrasonic pulses. Animal Behaviour, 10(3), 300–304. https://doi.org/10.1016/0003-3472(62)90053-2

Roeder, K. D. (1964). Aspects of the noctuid tympanic nerve response having significance in the avoidance of bats. Journal of Insect Physiology, 10(4), 529–546. https://doi.org/10.1016/0022-1910(64)90025-3

Roeder, K. D. (1967). Turning tendency of moths exposed to ultrasound while in stationary flight. Journal of Insect Physiology, 13(6), 873–888. https://doi.org/10.1016/0022-1910(67)90051-0

Rydell, J. (1998). Bat defence in lekking ghost swifts (*Hepialus humuli*), a moth without ultrasonic hearing. Proceedings of the Royal Society of London. Series B: Biological Sciences, 265(1404), 1373– 1376. https://doi.org/10.1098/rspb.1998.0444

Schmidt, S., Hanke, S., & Pillat, J. (2000). The role of echolocation in the hunting of terrestrial prey— New evidence for an underestimated strategy in the gleaning bat, *Megaderma lyra*. Journal of Comparative Physiology A: Sensory, Neural, and Behavioral Physiology, 186(10), 975–988. https://doi.org/10.1007/s003590000151

Schmitz, O. (2017). Predator and prey functional traits: Understanding the adaptive machinery driving predator–prey interactions. F1000Research, 6.

Schmitz, O. J., Buchkowski, R. W., Burghardt, K. T., & Donihue, C. M. (2015). Chapter Ten - Functional Traits and Trait-Mediated Interactions: Connecting Community-Level Interactions with Ecosystem Functioning. In S. Pawar, G. Woodward, & A. I. Dell (Eds.), Advances in Ecological Research, 52, 319–343. Academic Press. https://doi.org/10.1016/bs.aecr.2015.01.003

Schmitz, O. J., & Trussell, G. C. (2016). Multiple stressors, state-dependence and predation risk— foraging trade-offs: Toward a modern concept of trait-mediated indirect effects in communities and ecosystems. Current Opinion in Behavioral Sciences, 12, 6–11. https://doi.org/10.1016/j.cobeha.2016.08.003

Schneider, C. A., Rasband, W. S., & Eliceiri, K. W. (2012). NIH Image to ImageJ: 25 years of image analysis. Nature Methods, 9(7), Article 7. https://doi.org/10.1038/nmeth.2089

Schnitzler, H.U., Moss, C. F., & Denzinger, A. (2003). From spatial orientation to food acquisition in echolocating bats. Trends in Ecology & Evolution, 18(8), 386–394. https://doi.org/10.1016/S0169-5347(03)00185-X

Schoeman, M. C., & Jacobs, D. S. (2011). The relative influence of competition and prey defences on the trophic structure of animalivorous bat ensembles. Oecologia, 166(2), 493–506. https://doi.org/10.1007/s00442-010-1854-3

Scoble, M. J. (1992). The Lepidoptera: Form, function and diversity. *Oxford University Press*

Shi, J., Chen, F., & Keena, M. A. (2015). Differences in Wing Morphometrics of *Lymantria dispar* (Lepidoptera: Erebidae) Between Populations That Vary in Female Flight Capability. Annals of the Entomological Society of America, 108(4), 528–535. https://doi.org/10.1093/aesa/sav045

Shiel, C. B., McAney, C. M., & Fairley, J. S. (1991). Analysis of the diet of Natterer’s bat *Myotis nattereri* and the common long-eared bat *Plecotus auritus* in the West of Ireland. Journal of Zoology, 223(2), 299–305. https://doi.org/10.1111/j.1469-7998.1991.tb04766.x

Siemers, B. M., & Schnitzler, H.-U. (2000). Natterer’s bat (*Myotis nattereri* Kuhl, 1818) hawks for prey close to vegetation using echolocation signals of very broad bandwidth. Behavioral Ecology and Sociobiology, 47(6), 400–412. https://doi.org/10.1007/s002650050683

Spitz, J., Ridoux, V., & Brind’Amour, A. (2014). Let’s go beyond taxonomy in diet description: Testing a trait-based approach to prey–predator relationships. Journal of Animal Ecology, 83(5), 1137– 1148. https://doi.org/10.1111/1365-2656.12218

Stylman, M., Penz, C. M., & DeVries, P. (2020). Large Hind Wings Enhance Gliding Performance in Ground Effect in a Neotropical Butterfly (Lepidoptera: Nymphalidae). Annals of the Entomological Society of America, 113(1), 15–22. https://doi.org/10.1093/aesa/saz042

Subramanian, K. A., Sivaramakrishnan, K. G., & Gadgil, M. (2005). Impact of riparian land use on stream insects of Kudremukh National Park, Karnataka state, India. Journal of Insect Science, 5(1), 49. https://doi.org/10.1093/jis/5.1.49

Surlykke, A., Filskov, M., Fullard, J. H., & Forrest, E. (1999). Auditory Relationships to Size in Noctuid Moths: Bigger Is Better. Naturwissenschaften, 86(5), 238–241. https://doi.org/10.1007/s001140050607

ter Hofstede, H. M., Goerlitz, H. R., Ratcliffe, J. M., Holderied, M. W., & Surlykke, A. (2013). The simple ears of noctuoid moths are tuned to the calls of their sympatric bat community. Journal of Experimental Biology, 216(21), 3954–3962. https://doi.org/10.1242/jeb.093294

ter Hofstede, H. M., & Ratcliffe, J. M. (2016). Evolutionary escalation: The bat–moth arms race. Journal of Experimental Biology, 219(11), 1589–1602. https://doi.org/10.1242/jeb.086686

Thornhill, R., & Alcock, J. (2013). The Evolution of Insect Mating Systems. In The Evolution of Insect Mating Systems. *Harvard University Press*. https://doi.org/10.4159/harvard.9780674433960

Tyrell, K. (1990). *The ethology of the Malayan false vampire bat (Megaderma spasma), with special emphasis on auditory cues used in foraging* (PhD Thesis). University of Illinois at Urbana-Champaign.

Vesterinen, E. J., Ruokolainen, L., Wahlberg, N., Peña, C., Roslin, T., Laine, V. N., Vasko, V., Sääksjärvi, I. E., Norrdahl, K., & Lilley, T. M. (2016). What you need is what you eat? Prey selection by the bat *Myotis daubentonii*. Molecular Ecology, 25(7), 1581–1594. https://doi.org/10.1111/mec.13564

Violle, C., Navas, M.-L., Vile, D., Kazakou, E., Fortunel, C., Hummel, I., & Garnier, E. (2007). Let the concept of trait be functional! Oikos, 116(5), 882–892. https://doi.org/10.1111/j.0030-1299.2007.15559.x

Wagner, D. L., Grames, E. M., Forister, M. L., Berenbaum, M. R., & Stopak, D. (2021). Insect decline in the Anthropocene: Death by a thousand cuts. Proceedings of the National Academy of Sciences, 118(2), e2023989118. https://doi.org/10.1073/pnas.2023989118

Wang, X., & Müller, R. (2009). Pinna-rim skin folds narrow the sonar beam in the lesser false vampire bat (*Megaderma spasma*). The Journal of the Acoustical Society of America, 126(6), 3311–3318. https://doi.org/10.1121/1.3257210

Wasserthal, L.T. (1993). Swing-hovering combined with long tongue in hawkmoths, an antipredator adaptation during flower visits. Animal-plant interactions in tropical environments, 77-87.

Wray, A. K., Peery, M. Z., Jusino, M. A., Kochanski, J. M., Banik, M. T., Palmer, J. M., Lindner, D. L., & Gratton, C. (2021). Predator preferences shape the diets of arthropodivorous bats more than quantitative local prey abundance. Molecular Ecology, 30(3), 855–873. https://doi.org/10.1111/mec.15769

